# Cell Surface Differences within the Genus *Methanosarcina* Shape Interactions with the Extracellular Environment

**DOI:** 10.1101/2025.03.17.643819

**Authors:** Amelia-Elena Rotaru, Ghazaleh Gharib, Abdalluh Jabaley, Konstantinos Anestis, Rhitu Kotoky

## Abstract

*Methanosarcina* are metabolically versatile methanogenic archaea that can perform extracellular electron transfer (EET), with important ecological and biotechnological implications. These archaea are broadly classified into two types (Type I and Type II) based on their energy metabolism and are also differ in their aggregation-disaggregation behavior, cell surface properties, and electron transfer strategies.

Type I *Methanosarcina* typically form large multicellular aggregates within a methanochondroitin extracellular matrix, thrive in organic-rich environments, play a key role in anaerobic digestion during wastewater treatment and can perform EET. However, their mechanism of EET remains unresolved. In contrast, Type II *Methanosarcina* rely on multiheme c-type cytochromes for EET and are better adapted to low-organic, mineral-rich environments such as deep-sea sediments and aquifers, where they contribute to methane emissions.

Despite their significance, the molecular mechanisms behind EET in *Methanosarcina*— particularly for Type I—remain poorly understood. This review highlights what is known and what is unknown regarding the surface biology of *Methanosarcina*, their EET strategies, and biogeochemical and industrial roles, emphasizing the need for further research to unlock their full potential in sustainable methane management.

## 1. Introduction

*Methanosarcina* are methane-producing Archaea that play crucial roles in biotechnology and climate processes, impacting wastewater treatment, carbon capture, and greenhouse gas emissions. *Methanosarcina* are globally distributed, with high taxon prevalence in wastewater digesters, sediments, peatlands, paddy soils, and other agricultural lands. (1) They appear to thrive especially well in human-impacted environments, such as wastewater digesters and cultivated soils. **(Fig.1a)**

Their resilience in these altered ecosystems can be attributed to their ability to withstand various environmental stresses: they endure sudden pH fluctuation, high salt concentrations, elevated ammonia levels, desiccation, and even oxygen exposure (2–5). This resilience is further strengthened by their unique metabolic versatility. Unlike other methanogenic Archaea, *Methanosarcina* utilize a diverse array of substrates as electron donors, including: i) diffusible gases like hydrogen, ii) methylated compounds like methanol and methylamines, iii) volatile fatty acids like acetate, and iv) extracellular electrons obtained directly from insoluble surfaces in their environment, such as minerals, metals, electrode surfaces or electrogenic bacteria. (6, 7) **(Fig. 1b)** This latter mechanism, known as extracellular electron uptake, sets them apart from other methanogens.

**Figure 1.**
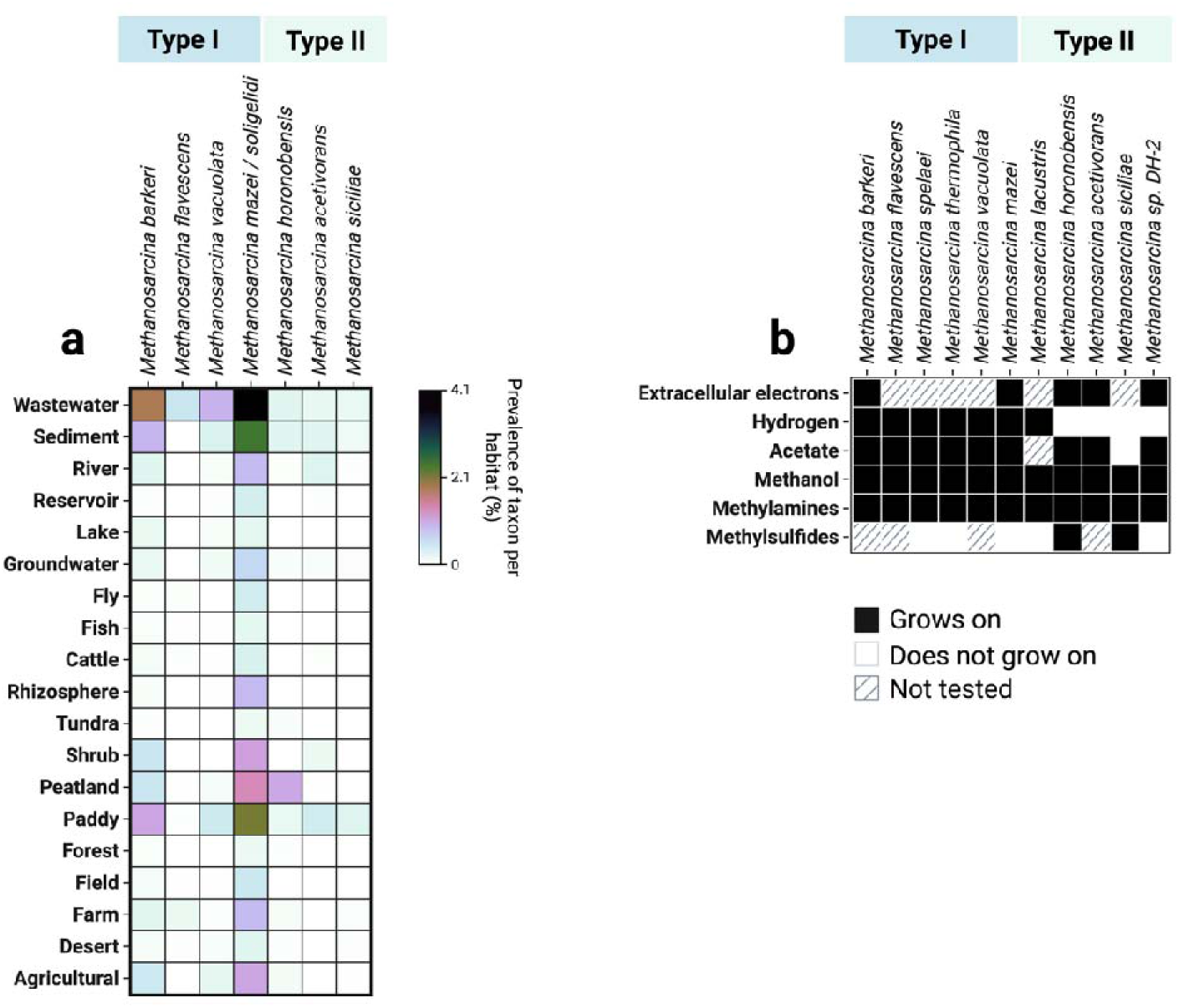
Habitats and growth phenotype for Type I and Type II *Methanosarcina.* (a) Distribution heatmap showing the prevalence (%) of different *Methanosarcina* species across various habitats, using taxon prevalence from microbeatlas.org. Color intensity corresponds to the relative abundance of each taxon, ranging from undetectable (white) to maximum prevalence (dark blue/black). (b) Growth phenotypes of various *Methanosarcina* species on six different substrates. Black squares indicate confirmed growth, white squares indicate absence of growth, and hatched squares denote untested conditions.

Additionally, *Methanosarcina* are genetically tractable and have been engineered to host synthetic pathways for producing biofuels (e.g., butanol) and other valuable compounds (e.g., terpene precursors like isoprene and lactate) (8, 9), thereby expanding their potential in biotechnological applications.

Their resilience to environmental stress, metabolic versatility, adaptability to human-impacted environments, genetic tractability, and capacity to harbor engineered pathways make *Methanosarcina* particularly valuable for research across environmental science, biotechnology and engineering fields (3, 6, 10).

In this review, we compare the two types of *Methanosarcina* (Type I and Type II) highlighting their distinct ecophysiologies, energy metabolisms, and cell surface properties, and exploring how these differences shape their interactions with the extracellular environment.

## 2. Habitat preferences

*Methanosarcina* species occupy a wide range of habitats, from natural aquatic sediments to engineered anaerobic digesters, where they engage in diverse ecological interactions with other organisms and surfaces in their environment. Originally, *Methanosarcina* were classified into two types based on their isolation sources: Type I strains – predominantly from freshwater or anaerobic digesters, and Type II strains – predominantly from marine environments (11). However, this classification has proven inadequate, and subsequent discoveries revealed both types in marine and freshwater habitats, as well as in wastewater, agricultural soils, rice paddies, and peatlands (**Fig. 1a**), indicating that salinity alone does not dictate their distribution. Instead, recent evidence suggests that the critical environmental factor is the organic matter content in their habitat (11).

Type I *Methanosarcina* prefer environments with high rates of organic matter degradation — such as anaerobic digesters and organic-rich sediments. These environments often have high concentrations of partially degraded plant material, supporting fermentative bacteria that release H_2_ as a waste product — an excellent electron donor for Type I *Methanosarcina* species — which can be used to reduce CO_2_ to methane. Their ability to rapidly utilize fermentation-derived substrates gives them a competitive advantage in organic-rich environments. (11, 12)

In contrast, Type II *Methanosarcina* prefer environments with lower organic content, including subsurface aquifers, sandy sediments and soils. In these habitats, Type II *Methanosarcina* may engage in alternative respiratory metabolisms including respiration of ferric-iron (Fe^III^) or oxidized humic substances (13–15), which likely confer a competitive advantage to Type II strains over Type I strains under these conditions.

## 3. Two physiological types of *Methanosarcina*

The most significant physiological difference between Type I and Type II *Methanosarcina* is their ability to use H_2_ as an electron donor: Type I can use H_2_, whereas Type II cannot. The difference is tied to variations in their electron transport chain and overall energy metabolism. These contrasts become especially apparent when examining acetate metabolism in representative strains such as *M. barkeri* (Type I) and *M. acetivorans* (Type II) **(Fig. 2).**

**Figure 2.**
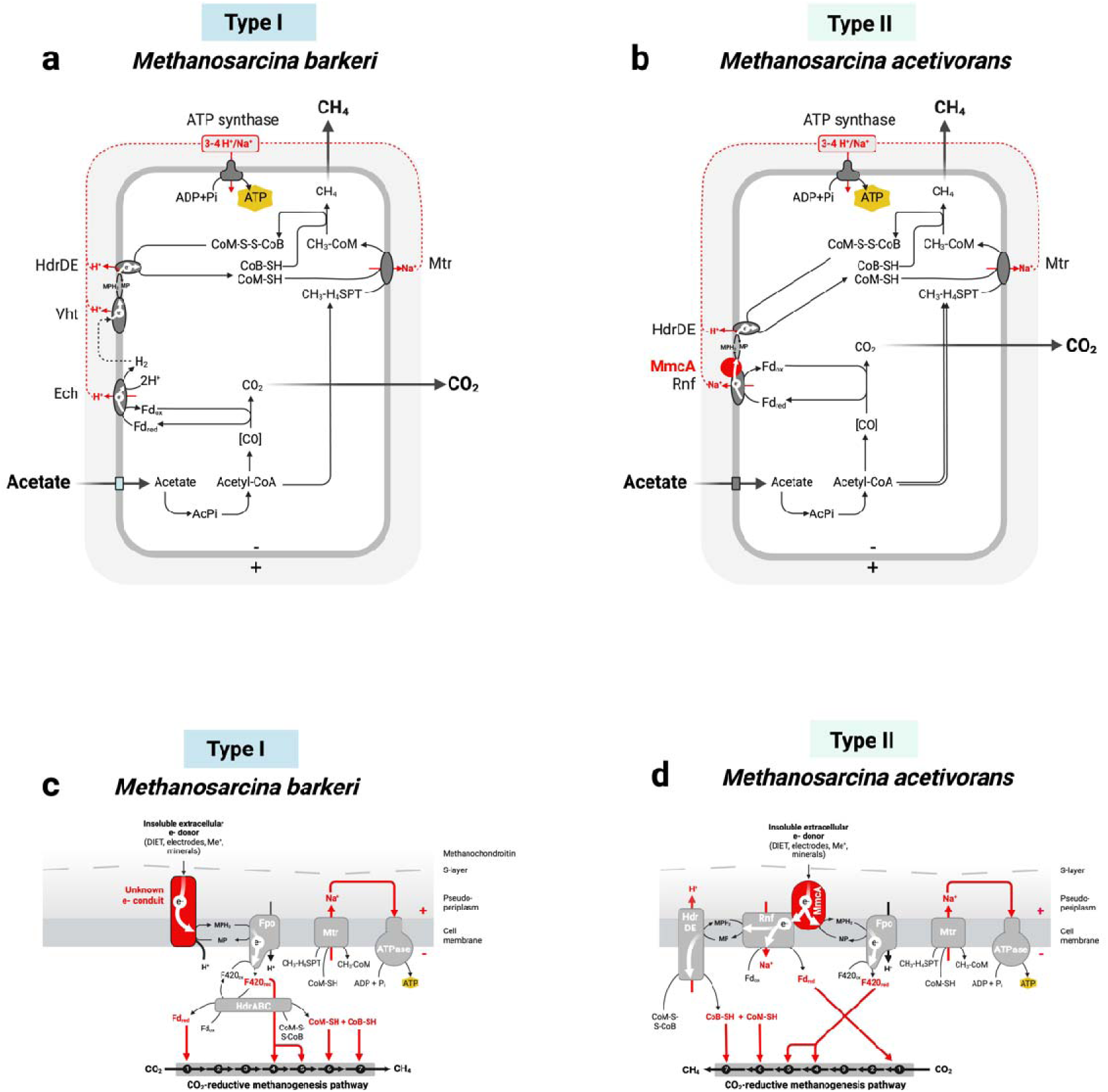
Schematic representation of energy conservation during acetoclastic methanogenesis (a, b) and CO_2_-reductive methanogenesis driven by extracellular electrons (c, d) in *Methanosarcina barkeri* (Type I) and *Methanosarcina acetivorans* (Type II). Panels (a, b) illustrate acetate uptake and conversion to methane and carbon dioxide, emphasizing the different membrane complexes. Panels (c, d) depict a proposed mechanism for accepting extracellular electrons directly – an alternative pathway for methanogenesis. Reactions 1-7 of CO_2_-reductive methanogenesis include: (1) CO_2_-activation by methanofuran, (2) formyl transfer to tetrahydromethanopterin (H_4_MPT), (3) cyclization to methenyl-H_4_MPT, (4) reduction to methylene-H4MPT, (5) further reduction to methyl-H4MPT, (6) methyl transfer to coenzyme M, (7) final reduction of methyl-coenzyme M to methane.

During acetate metabolism, *M. barkeri* (Type I) employs an Ech (Energy Converting Hydrogenase) complex to generate H_2_ in the cytoplasm on account of acetate-derived reduced ferredoxin while simultaneously pumping protons into the periplasm. The cytoplasmic H_2_ then diffuses across the membrane and is immediately recycled by the Vht dehydrogenase (H_2_-cycling), which transfers electrons to methanophenazine, while simultaneously pumping protons into the periplasm. Reduced methanophenazine subsequently donates electrons to the terminal complex in this electron transport chain - the membrane-bound HdrDE (heterodisulfide reductase), facilitating the reduction of coenzyme M (CoM) and coenzyme B (CoB) and contributing to increasing the proton motive force (16, 17).

In contrast, Type II – *M. acetivorans* lacks H_2_-cycling and instead relies on the Rnf-complex to conserve energy from acetate metabolism. Reduced ferredoxin derived from acetate donates electrons to the Rnf-complex, while contributing to ion translocation across the membrane. Electrons then flow from the Rnf to the methanophenazine pool via a membrane-bound multiheme c-type cytochrome (MmcA) present in all Type II *Methanosarcina* but absent in Type I. Electrons from methanophenazine reach the HdrDE complex, fueling CoM-CoB reduction while also contributing to ion translocation increasing the ion motive force (15, 18).

Ultimately, the built-in ion-motive in both Type I and Type II *Methanosarcina* fuels ATP synthesis via a promiscuous ATP synthase concurrently coupled to both Na^+^ and H^+^ translocation (19).

## 4. Interactions with the extracellular environment

*Methanosarcina* establish co-dependent metabolic associations with certain bacteria to overcome resource limitations and energetic challenges that neither organism could manage alone. In these syntrophic associations, the bacterial partners must dispose of excess reducing equivalents in the methanogenic zone where the only remaining electron acceptor is CO_2_. If they fail to do so, respiratory bacterial partners cannot carry out their oxidative metabolism with CO_2_ is the terminal electron acceptor. Comparably, fermentative partners face feedback inhibition from the buildup of reduced waste products (e.g., H_2_). On the other hand, *Methanosarcina* cannot independently oxidize complex organics and instead depend on the reduced compounds provided by the bacterial partners, such as H_2_ (or electrons directly) to reduce CO_2_ to methane. This mutually beneficial syntrophic relationship hinges on the bacterial partner carrying out a thermodynamically unfavorable oxidation step, made feasible by the methanogen’s rapid consumption of electrons (6).

Once released by the bacterial partner, electrons can reach *Methanosarcina* via:

i) Hydrogen Interspecies Transfer (HIT) – relying on diffusible H_2_ produced by the partner bacteria (e.g., *Desulfovibrio vulgaris*) (20)
ii) Direct Interspecies Electron Transfer (DIET) – relying on a network of electrically conductive cell surface structures that enable direct electron transfer. (e.g., *Geobacter metallireducens*) (21)
iii) Conductive particle – mediated Interspecies Electron Transfer (CIET) – relying on minerals (e.g., magnetite)(22), or other conductive particles (e.g., activated carbon, biochar)(23, 24) that act as electron-transfering “bridges” between partner bacteria and *Methanosarcina*, diminishing the need of the two partners to produce their own EET machinery (25).

A unique feature of all *Methanosarcinales* that differentiates them from other methanogens is their ability to uptake electrons from the extracellular environment (e.g., from conductive particles, or other cells). Both Type I and Type II *Methanosarcina* can engage in syntrophic interactions with bacteria capable of extracellular electron transfer (EET) such as *Geobacter* species (11, 21, 26–28) or *Rhodoferax* (28).

However, only Type I *Methanosarcina* have been observed forming partnerships reliant on interspecies H_2_ transfer (e.g., with *Pelobacter carbinolicus* (21), *Smithella propionica*, *Syntrophobacter wollinii* (29) or *Desulfovibrio vulgaris* (20). Additionally, Type I *Methanosarcina* can receive electrons directly from cathodes (at ∼-400 vs. the standard hydrogen electrode), and from syntrophic partners indirectly via conductive particles (22, 23, 28). However, the specific mechanism enabling electron uptake in *Methanosarcina* Type I remains unclear (**Fig. 2c**). A previous study suggested that electrons from extracellular electron donors enter the cell via an unknown cell surface component that is redox-active, and then enter the methanophenazine pool, subsequently reaching the membrane-bound Fpo-complex, which likely operates in reverse to reduce F_420_. The reduced F_420_ could then drive the intra-cytoplasmic reduction of ferredoxin and CoM-S-S-CoB generating all the reduced carriers to complete the CO_2_-reductive methanogenesis steps (30).

In contrast, Type II *Methanosarcina* (*M. acetivorans, M. horonobensis)* primarily receive electrons from other cells directly or via conductive particles (31, 32). Although some studies showed that *M. acetivorans* can receive extracellular electrons from Fe^0^ when provided with minimal acetate (*M. acetivorans*), these findings were not consistently reproduced by others or in other Type II strains (*M. horonobensis*) (33). Intriguingly, *M. horonobensis*, could also not accept electrons from a cathode after repeated tests (31). It remains unclear why this is the case. Notably, *M. acetivorans* requires a membrane-bound multiheme c-type cytochrome (MmcA) for effective EET to and from redox-active extracellular surfaces (**Fig. 2d**). This cytochrome is bidirectional and crucial for both intracellular and extracellular electron flow (15). Its absence renders *M. acetivorans* incapable of EET, including DIET (13) or reduction of extracellular electron acceptors (13, 15). Thus, the EET mechanism in *M. acetivorans* mirrors that of electroactive bacteria employing membrane-bound multiheme cytochromes (34). Interestingly, the distribution of *mmcA*-gene homologs extends beyond *Methanosarcina* Type II strains, to marine species within the family *Methanosarcinaceaea*, including *Methanococcoides*, *Methanohalophylus*, *Methanosalsum* and *Methanolobus* suggesting they may have the ability to perform EET (**Fig. 3a**) (15).

**Figure 3.**
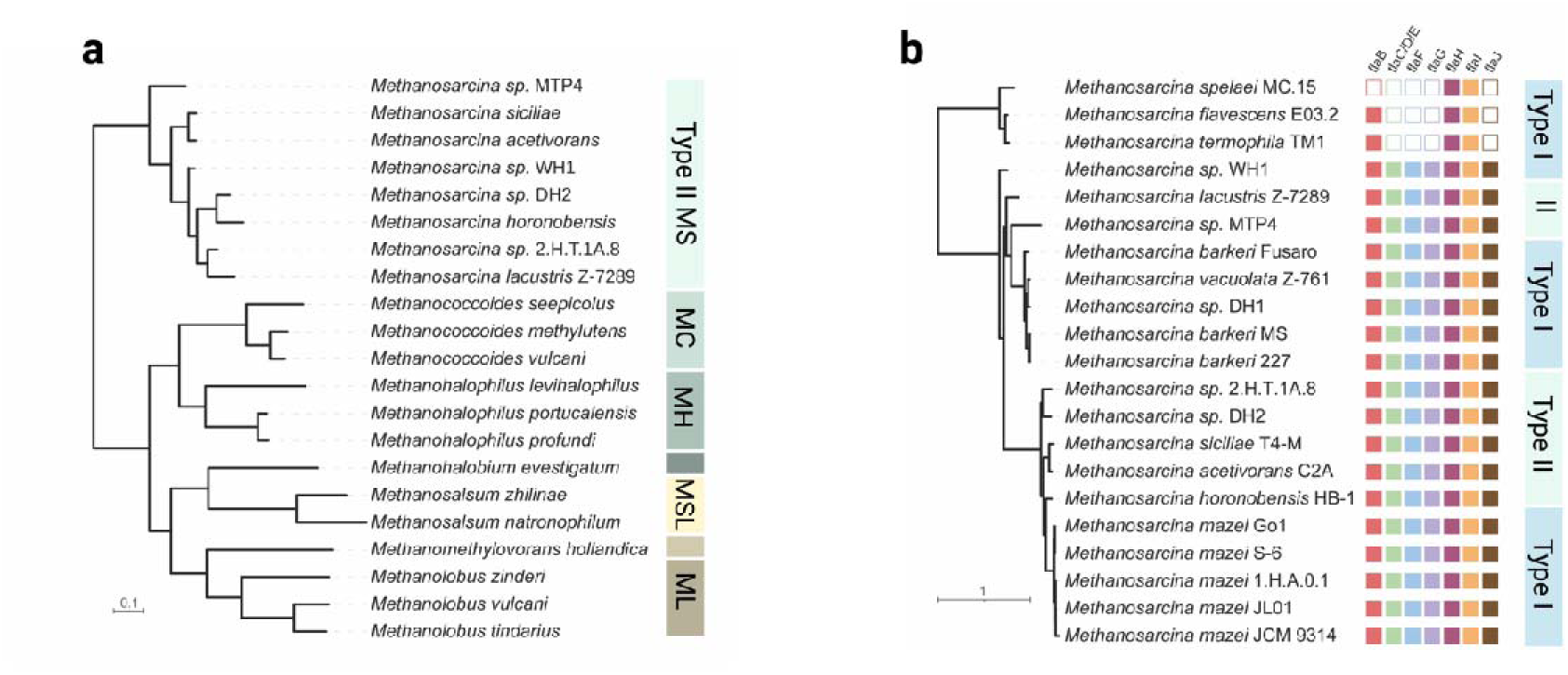
Comparative phylogeny of. (a) the *mmcA*-gene and (b) the *fla*-operon. (a) Phylogenetic relationships of the multiheme c-type cytochrome genes (*mmcA*) in Type II *Methanosarcina* and their gene orthologs in other *Methanosarcinaceae*. Colored blocks group the distinct genera abbreviated MS for *Methanosarcina*, MC for *Methanococcoides*, MH for *Methanohalophilus*, MSL for *Methanosalsum*, and ML for *Methanolobus*. (b) Distribution of the *fla*-operon among Type I and Type II *Methanosarcina*. Filled squares indicate presence or empty squares indicate absence of specific archaellum-related genes. Scale bar indicates amino acid substitutions per site.

Below, we examine how Type I *Methanosarcina* differ in terms of cell surface properties and explore how these differences may lead to a distinct mechanism for extracellular electron transfer – one that deviates from the conventional MHC-dependent EET pathways.

## 5. Distinct cell surfaces for Type I and Type II *Methanosarcina*

Diving into the cell from the exterior, we first encounter turfs of flagella-like structures (archaella) and a unique extracellular polysaccharide layer called methanochondroitin, along with extracellular enzymes that break down this polymer. Beneath these components lies a proteinaceous S-layer encasing the periplasm. Interestingly, in these methanogens, the periplasm is uniquely bound by a diether-lipid membrane (archaeol) on the cytoplasmic side and the S-layer on the extracellular side. In Type II species, this inner membrane also houses the MmcA multiheme *c*-type cytochrome crucial for extracellular electron transfer.

### 5.1. Archaella

Cell motility in *Methanosarcina* is facilitated by a cell surface structure found in many archaea – the archaella – that likely emerged by convergent evolution in all kingdoms of life. The archaella shares functional homology to bacterial flagella and eukaryotic cilia but is distinct in its assembly and operation (35, 36). The term ‘archaellum’ was introduced by Jarrell and Albers in 2012 (37) to highlight the uniqueness of the archaeal motility apparatus compared to its bacterial and eukaryotic counterparts. Although the archaellum shares certain similarities with the bacterial type IV pili system - such as the involvement of FlaK/PibD peptidases in the posttranslational modification of its subunits and the presence of homologs of *flaI* and *flaJ*-genes (38), several genes (*flaCDEFGH*) remain unique to Archaea. Knockout studies have confirmed the essential role of these genes, as their deletion results in non-functional archaella (39–42). Within the *Methanosarcina* genus, the genetic toolkit for archaellum biosynthesis is widely conserved across both Type I and Type II strains, does not follow the Type I/Type II delineation, and only a few Type I species (*M. spelaei*, *M. thermophila* and *M. flavescens*) lack entire gene clusters (**Fig. 3b**).

Beyond motility, the archaellum of *Methanosarcina* species may facilitate DIET syntrophic relationships by serving as an electron conduit (32). Long-range electron transport via archaella is thought to be facilitated by its high abundance of aromatic residues, such as phenylalanine (43–45). This dual role in electron transport and motility is similar in bacterial type IV pili, which serve as electrically conductive e-pili, enabling DIET and EET to insoluble substrates (25, 46–48).

Recent studies have extended these observations to Archaea; for instance, the archaellum of the methanogen *Methanospirillum hungatei*, has been shown to exhibit electrical conductivity. In *Methanosarcina acetivorans*, deletion of archaellin-encoding genes encoding inhibited electron exchange via DIET with *Geobacter metallireducens*, an effect that could be compensated by the addition of electrically conductive granular activated carbon (GAC) (13). Interestingly, archaellum gene expression appears consistent under both DIET and monoculture conditions, suggesting constitutive expression. However, it remains unclear whether the observed impact on DIET arises from the archaellum’s conductive properties or its roles in motility, attachment, and partner location (49).

In summary, while the archaellum is essential for *Methanosarcina’s* motility, it’s other role in electron transfer merits further investigation to delineate its contribution to both physical and metabolic interactions within syntrophic communities.

### 5.2. Methanochondroitin

*Methanosarcina* must interact effectively with its environment and in many species the first point of contact is a unique heteropolysaccharide called methanochondroitin – a polymer whose chemical structure resembles the chondroitin sulfate in mammalian connective tissues (50, 51). Unlike its mammalian counterpart, methanochondroitin is not sulfated; it rather consists of repeating trimers of two N-acetylglucosamine units and one glucuronic acid in the configuration [→4)-β-D-GlcUA(1→3)-D-GalNAc-(1→ 4)-D-GalNAc(1 →]_n_ (52).

Although, a biosynthetic pathway for this polymer was suggested 30 years ago, based on isolated intermediates (53), neither the specific mechanisms of biosynthesis nor the genes involved have been identified.

Methanochondroitin is not a static barrier, but a dynamic layer that responds to environmental cues. Its presence or absence can be modulated by changes in salinity (54) or osmolarity (55) thereby influencing how *Methanosarcina* interacts with its environment. This layer is observed in Type I *Methanosarcina* such as *M. barkeri*, which prefer low-osmolarity environments. Under these conditions, they can produce a substantial methanochondroitin layer. Biophysical measurements indicate a total thickness of up to 170 nm around each cell, accounting for anything between ∼20%-50% of the cellular volume. By contrast, Type II species, like *M. acetivorans*, which thrive in high-osmolarity marine habitats, often show a reduced or absent methanochondroitin layer relying primarily on their S-layer for external protection. In **Fig. 4**, this contrast is evident when *M. barkeri* (Type I) grown in freshwater media - exhibits a robust methanochondroitin layer encasing the cell membranes whereas, a *M. acetivorans* (Type II) grown in marine medium lacks it.

**Figure 4.**
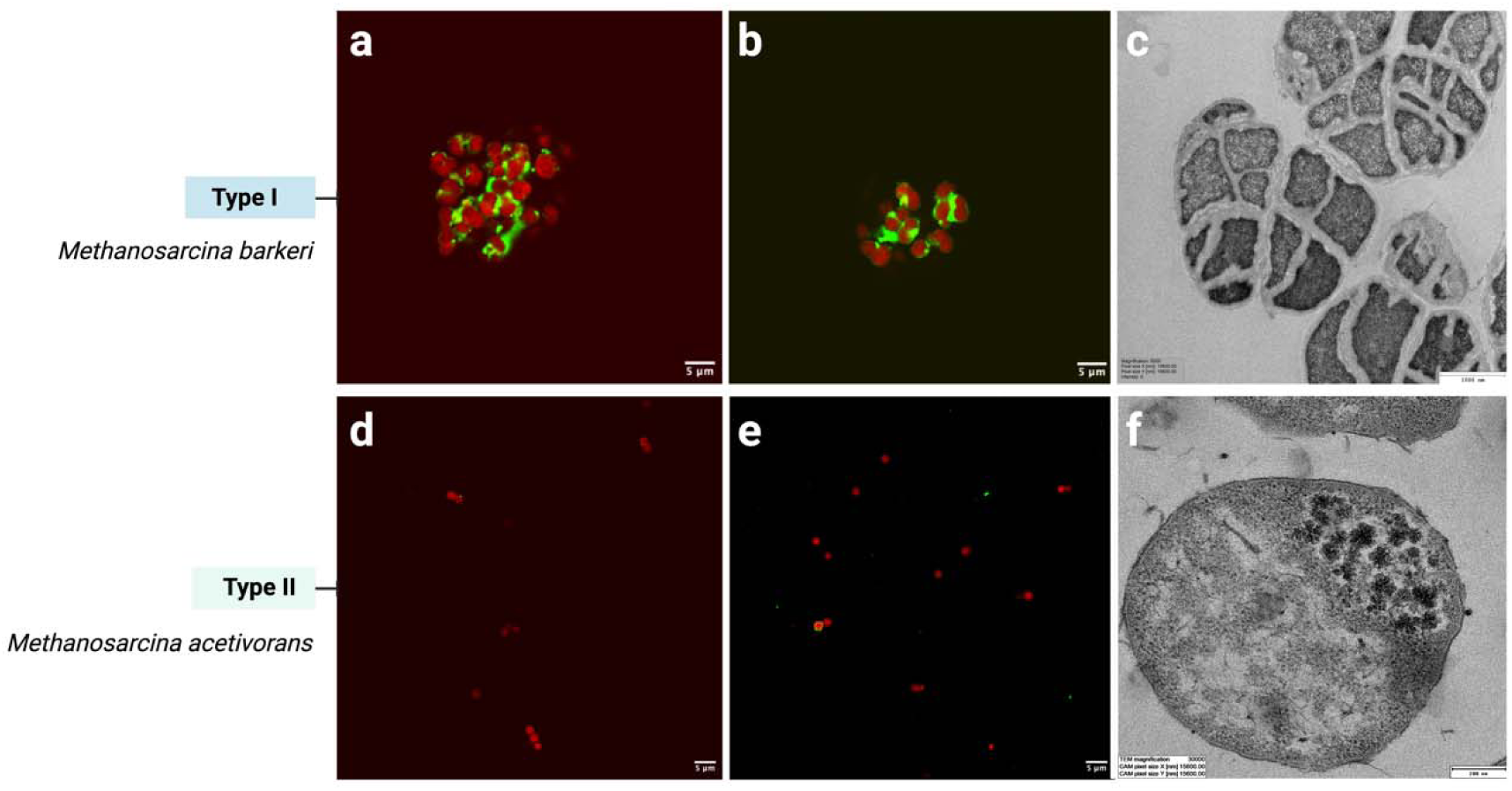
Micrographs of Type I – *Methanosarcina* barkeri (a-c) and Type II *Methanosarcina acetivorans* (d-f). (a, b) Fluorescence images of *M. barkeri* stained with a membrane dye (red) and concanavalin A (green) targeting the extracellular methanochondroitin layer. (c) Transmission electron micrograph (TEM) of *M. barkeri*, showing densely packed cells in aggregates enveloped in a thick extracellular methanochondroitin layer (scale bar = 200Lnm). In contrast, *M. acetivorans* typically grows as individual cells and lacks a visible methanochondroitin coat (d, e). Fluorescence images by Abdalluh Jabaley. TEM images by Pia Bomhold Jensen and Abdalluh Jabaley.

The formation of a methanochondroitin layer in low osmolarity environments also influences cell division. Daughter cells remain embedded within the existing matrix – rather than fully separating – leading to multicellular aggregates. Conversely, in higher osmolarity environments, cells propagate individually surrounded only by the S-layer (56). Remarkably, both Type I and Type II *Methanosarcina* can grow in both low and high osmolarity environments (54), transitioning between solitary and aggregate states as needed.

Functionally, methanochondroitin appears to support *Methanosarcina* under various stresses providing structural cohesion and protection analogous to Gram positive bacterial cell walls. It effectively “glues” the cells together in a cluster shielding the cells in the interior of the aggregate from external stressors. For example, in *M. mazei* cells in the interior of multicellular aggregates resist infection by *Methanosarcina* spherical virus, which targets the S-Layer (57). Similarly, methanochondroitin-related aggregation has been tied to heightened survival under oxygen stress (58), desiccation, or elevated temperatures (2). In addition, the negatively charged methanochondroitin layer can sequester toxic metal ions such as cadmium, thus mitigating heavy metal toxicity (59).

An intriguing possibility is that methanochondroitin contributes to extracellular electron transfer. Chondroitin sulfate in mammalian systems is known to conduct electric current (60), raising the prospect that methanochondroitin could have similar properties. Such conductivity may mirror mechanism observed in electroactive bacteria such as *Geobacter sulfurreducens,* where extracellular polysaccharides facilitate electron flow by either forming conductive matrices (61) possibly by sequestering electroactive atoms (e.g. Fe) or molecules (e.g., multiheme c-type cytochromes (62).

Thus, methanochondroitin extends well beyond a structural layer. It underlies multicellularity, influences environmental adaptation, roles in cellular aggregation, and may even contribute to electron transfer processes with its presence and thickness varying between Type I and Type II as an adaptive response to their respective ecological niches.

### 5.3. Extracellular disaggregating enzyme

Some *Methanosarcina* strains transition between aggregated (multicellular) and dispersed (unicellular) states as part of their life cycle (63, 64). For instance, in *M. mazei* this transition is controlled by an extracellular enzyme known as disaggregatase. This enzyme specifically targets only β-1,4-glycosidic bonds between glucuronic acid and galacturonic acid in methanochondroitin, an extracellular polysaccharide unique to *Methanosarcina* (65).

Currently, the disaggregatase has only been characterized in strains of *M. mazei* (Type I) (66). Early observations suggested the existence of this enzyme in a strain (*M. mazei* LYC) which spontaneously dispersed during growth (65). During the initial growth stages, LYC’s single cells divide but remain partially attached and encased in the methanochondroitin matrix, building large aggregates. Upon reaching exponential growth, the aggregates begin secreting the disaggregatase, causing the surrounding methanochondroitin matrix to degrade and the large clumps of cells to dissassemble into individual coccoid cells. At this point, the culture medium becomes turbid as aggregates break apart (54). This enzymatic dispersal is sometimes effective on the producer strain but sometimes on other *M. mazei* strains or even other *Methanosarcina* species (e.g., *M. thermophila*) (65, 67).

Although the enzymatic activity and extracellular localization of the disaggregatase has been experimentally confirmed only in *M. mazei*, genomic screenings have identified dissagregatase-related domains in other *Methanosarcina* species, including another Type I (*M. barkeri*) and a Type II species (*M. acetivorans*). However, neither *M. barkeri* or *M. acetivorans* cells interact with antibodies targeting the *M. mazei* disagregatase, suggesting that their dissaggregatase may either be absent or sufficiently different (67). It has been hypothesized that these other species may rely on environmental cues rather than enzymatic action to trigger aggregate dispersal, although this hypothesis awaits experimental confirmation (55).

The timing of aggregate dispersal can significantly impact the ecological fitnes of *Methanosarcina*. Aggregation protects the cells under adverse environmental conditions, while timely disaggregation when resources fluctuate or are limited may promote nutrient uptake, or access to new ecological niches. This enzymatic transition between multicellular and unicellular states could serve as an adaptive mechanism, balancing the need for protection with the benefits of mobility (64, 68).

In addition to its role in dispersal, the secretion of disaggregatase may influence microbial community dynamics in mixed biofilms. It could disrupt competitors, or alter community structure in favor of disaggregatase-producing *Methansoarcina* strains. This enzymatic strategy might provide a competive advantage, enhancing nutrient access and persistence in diverse microbial ecosystems (69, 70).

Finally, the role of the dissaggragatase in biofilm matrix disassembly raises intriguing questions about its potential impact on extracellular electron transfer (EET). If the methanochondroitin matrix supports EET activity, its enzymatic breakdown might act as an “off switch” for cellular electroactivity, possibly linking multicellular organization with energy metabolism strategies.

### 5.4. S-layer

All *Methanosarcina* species, both Type I and Type II, are surounded by a proteinaceous surface layer (S-layer) (71) composed predominantly of a single glycosylated protein (100-130 kDa), arranged in a porous hexagonal lattice (72, 73). Structural analyses of the S-layer protein from *M. acetivorans* (Type II) revealed four characteristic regions: (i) an N-terminal signal peptide, (ii) tandem-duplicated DUF1608 domains, (iii) a negatively charged tether (∼60 amino acids), and (iv) a C-terminal transmembrane helix that possibly anchors the S-layer to the cytoplasmic membrane (74). High conservation of the major S-layer protein across *Methanosarcina* species (including Type I species - *M. barkeri*, *M. mazei* and Type II species - *M. acetivorans*) led to the classification of a new protein family - the *Methanosarcinale* S-layer Tile Protein (MSTP) family characterized by DUF1608 domains (71). Additional support for this protein family came from phylogenetic analsyes, which confirmed DUF1608-domain containing proteins in all sequenced genomes of *Methanosarcina* (75) (**Fig. 5a**). No structural differences in latice symetry or organization have been reported between Type I and Type II *Methanosarcina* S-layer proteins. Fascinatingly, the S-layer is considered one of the most primitive cellular envelope structures, predating the divergence of major archaean lineages. Comparison of S-layer protein sequence of *Methanosarcina* sps. with other groups of Archaea showed that there is significant sequence conservation among the Methanosarcina species and forms a compact cluster (**Fig. 5b**), suggesting a common S-layer architecture for Methanosarcinaceae. For other groups as well, the s-layer protein, has high sequence identity within the same species but different groups (*Thermococcus*, *Methanococcus*, *Halobacterium*) formed separate clusters, suggesting evolutionary differences in different archaean groups. It is though to have originated before the evolution of the murein-containing cell envelope/sacculus. The *Methanosarcina* S-layer protein has a β-sandwich domain structurally homologous to eukaryotic RNA virus-coat proteins, hinting that early cells and viruses convergently evolved hard protein latices for protection (74).

**Figure 5.**
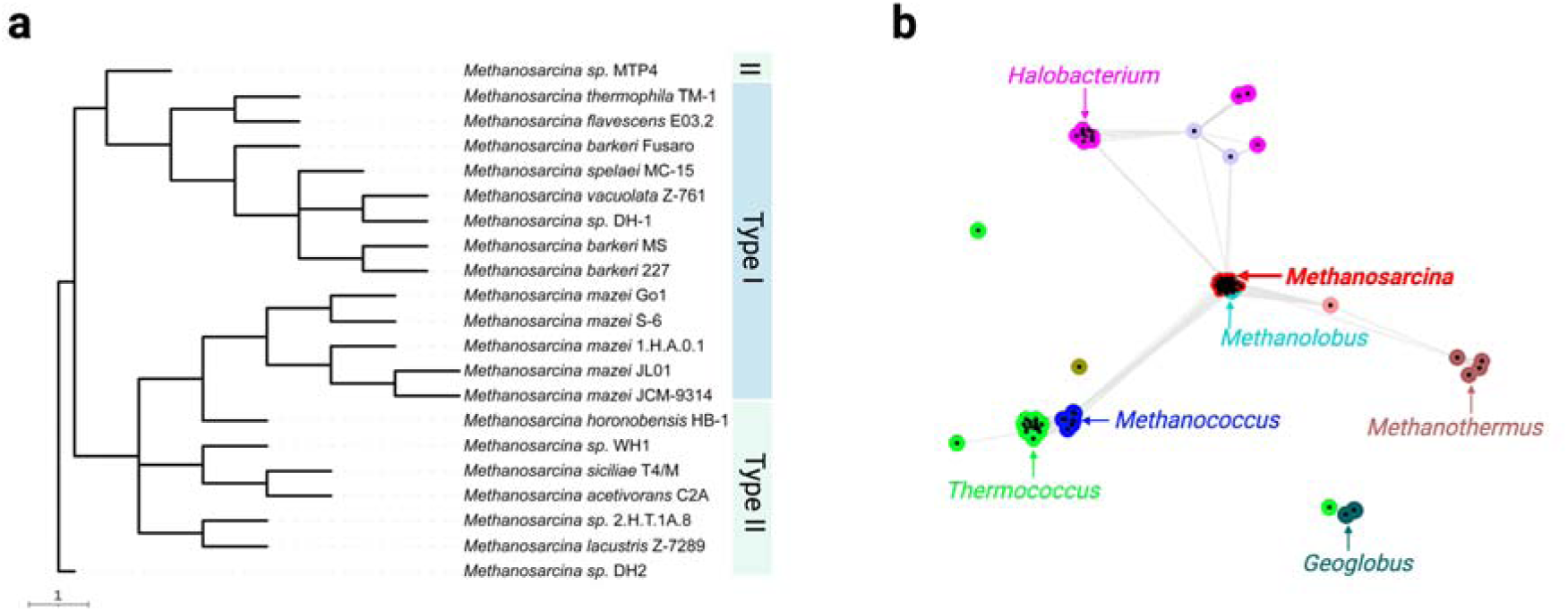
Phylogenetic and network analysis of S-layer protein orthologues. (a) Phylogenetic tree of selected *Methanosarcina* S-layer proteins constructed by Neighbor-Joining in MEGA11. The tree highlights the two types of *Methanosarcina* (Type I – light blue and Type II – light green). Scale bar represents substitutions per nucleotide site. (b) Network visualization based on amino acid sequence relatedness of archaeal S-layer proteins clustering of different archaeal groups: *Methanosarcina* (red), *Methanolobus* (cyan), *Methanococcus* (dark blue), *Thermococcus* (bright green), *Geoglobus* (dark green), *Halobacterium* (pink). Dots represent individual surface-layer proteins, and a high sequence similarity (blast p values) is represented by connecting lines.

Although the precise functions of the archaeal S-layer remain debated, it is generally accepted to serve as a protective coat and molecular sieve, mediating nutrient uptake, waste excretion, surface recognition, cellular signaling, defensive interactions with their environment (76, 77). In Type II *Methanosarcina*, the S-layer represents the primary protective barier surounding the lipid membrane, whereas Type I *Methanosarcina* leverage both the S-layer and an additional methanochondroitin layer for environmental responses and protection (See 5.2).

Interestingly, transcriptomic studies revealed increased expression of S-layer proteins in both Type I (*M. barkeri*) and Type II (*M. acetivorans*) *Methanosarcina* grown via DIET (Direct interspecies electron transfer) with an electrogenic syntroph, compared to growth on soluble (acetate) or diffusible substrates (H_2_ from a partner hydrogen-generating syntroph). This observation suggests a potential role of the S-layer in electron transfer processes possibly by docking electron-carrying molecules to create an electron-conductive interface at the cell surface. For instance, in *M. acetivorans* (Type II) the S-layer could facilitate electron transfer by docking the multiheme cytochrome responsible for EET in this organism (MmcA). However, in Type I *Methanosarcina,* which lack such cytochromes, the exact mechanism and involvment of the S-layer remains unresolved.

## 6. Implications

*Methanosarcina* are versatile methanogenic Archaea, adept at colonizing diverse habitats with their broad substrate utilization and flexible energy metabolism. Their unique cell surface properties help them withstand environmental stressors. Based on the energy-conserving complexes they harbor, *Methanosarcina* are classified into Type I (Ech-dependent) and a Type II (Rnf-dependent), each exhibiting unique cell surface compositions promoting habitat-specific survival and propagation, and possibly linked to distinct strategies to perform extracellular electron transfer. This distinction between Type I and Type II *Methanosarcina* is functionally relevant, shaping their survival strategies, interactions and contribution to methane cycling across diverse ecosystems.

Type I *Methanosarcina* form multicellular aggregates encased in methanochondroitin, which provides an extra protective barrier in addition to the glycosylated S-layer shared by all *Methanosarcina*. Type I cells also produce disaggregatase, an enzyme that controls methanochondroitin breakdown and aggregate dispersal. These *Methanosarcina* also engage in DIET and CIET syntrophy and can extract electrons from poised cathodes (Table 1) without relying on typical EET molecules – multiheme *c*-type cytochromes (MmcA). They may instead rely on their methanochondroitin layer to sequester redox-active required for EET. These type I *Methanosarcina* often govern anaerobic digesters (AD) and rice paddies where organic loads and pollutants are high. Multiple studies have shown that *Methanosarcina* operating in CIET partnerships can accelerate the conversion of organics from digestate to methane, a promissing strategy for the wastewater treatment industry. Indeed promotion of AD with conductive particles promoting CIET-partnerships has reached pilot-scale trials (78, 79). Furthermore, augmenting CIET-partners on conductive support further benefits the process (80). Given their proven capacity of DIET and CIET, and ability to withstand high loads of organic and pollutants, Type I *Methanosarcina* are expected to play pivotal roles in emerging bioelectrochemical approaches designed to assist AD and remediate industrial waters.

**Table 1.**
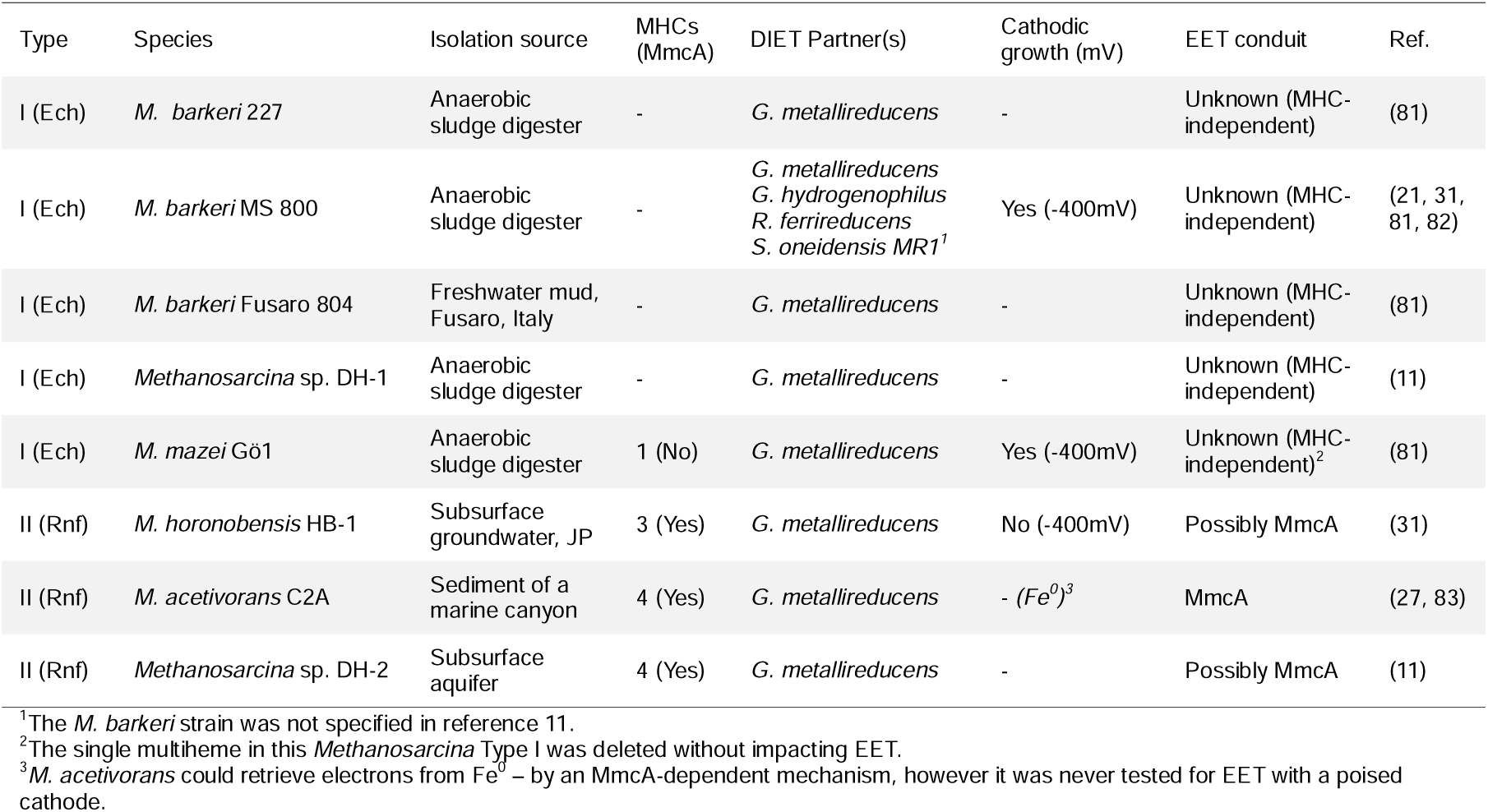
Overview of representative Type I and Type II *Methanosarcina* species evaluated for DIET syntrophy with electroactive bacteria and for cathodic growth under conditions of minimal electrochemical H□ evolution.

By contrast, Type II *Methanosarcina* generally grow as single cells without the additional methanochondroitin layer or the need for enzymatic dispersion; solely covered by the glycosylated S-layer which likley anchors the multiheme cytochrome MmcA that is crucial for EET. This MmcA supports both respiratory metabolism with Fe(III)-minerals, and electron uptake via DIET from *Geobacter*, or from Fe^0^ (Table 1). Although less prevalent in the environment, Type II lineages can displace Type I under conditions of low organic load and limited nutrients, such as deep marine sediments or deep subsurface aquifers. In these environments, the availability of mineral-based electron acceptors and suitable DIET partners offers essential redox-active substrates, allowing Type II *Methanosarcina* to access otherwise inaccessible energy sources to produce methane. These Type II *Methanosarcina* may be especially important in green house gas emission from marine and lacustrine environments.

In conclusion, the differences between Type I and Type II *Methanosarcina* extend beyond energy metabolism to fundamental differences in aggregation-disagregation, cell surface properties and extracellular electron transfer strategies shaping their ecological roles. Type I *Methanosarcina*, typically form multicellular aggregates encased in a methanochondroitin layer, dominate organic rich environments such as anaerobic digesters. In contrast, Type II *Methanosarcina* rely on multiheme *c*-type cytochromes for electron transfer, live as single cells, in low-organic, mineral-rich environments (deep sea sediment and aquifers), where they contribute to methane emissions.

*Methanosarcina* have unique EET capabilities between methanogenic Archaea, fundamental questions remain, particularly regarding the mechanism of EET in Type I species, which lack the known cytochrome-based system found in Type II. However, the differences between type I and type II suggests that these organisms have convergently evolved the ability for EET likely driven by similar evolutionary constraints in their habitats. Delving deeper into their electron transfer pathways and exploring their interactions with other microorganisms will be crucial for uncovering their full ecological importance, understanding their contribution to global carbon cycling, and assessing their potential for future industrial uses.

## Aknowledgements

This article is a contribution to a Novo Nordisk Ascending Investigator grant NNF21OC0067353 and an ERC Consolidator grant awarded to AER. We would like to thank the Danish Molecular Biomedical Imaging Center, at SDU for access and training on their confocal microscope, and Thomas Boesen and Pia Bomholt Jensen for access and training at the Cryo-EM facility at Aarhus University. Figures were compiled using Biorender.

## Notes

### Competing Interest Statement

The authors have declared no competing interest.

